# Discovery of inhibitors for bacterial Arr enzymes ADP-ribosylating and inactivating rifamycin antibiotics

**DOI:** 10.1101/2025.02.20.639278

**Authors:** Juho Alaviuhkola, Sondos Abdulmajeed, Sven. T. Sowa, Lari Lehtiö

**Affiliations:** Faculty of Biochemistry and Molecular Medicine & Biocenter Oulu, University of Oulu, Finland

## Abstract

ADP-ribosylation is an enzymatic process where an ADP-ribose moiety is transferred from NAD^+^ to an acceptor molecule. While ADP-ribosylation is well-established as a post-translational modification of proteins, rifamycin antibiotics are its only known small-molecule targets. ADP-ribosylation of rifampicin was first identified in *Mycolicibacterium smegmatis,* whose Arr enzyme transfers the ADP-ribose moiety to the 23-hydroxy group of rifampicin preventing its interaction with the bacterial RNA polymerase thereby inactivating the antibiotic. Arr homologues are widely spread among bacterial species and present in several pathogenic species often associated with mobile genetic elements. Inhibition of Arr enzymes offers a promising strategy to overcome ADP-ribosylation mediated rifamycin resistance. We developed a high-throughput activity assay, which was applied to screen an in-house library of human ADP-ribosyltransferase-targeted compounds. We identified 15 inhibitors with IC_50_ values below 5 µM against four Arr enzymes from *M. smegmatis*, *Pseudomonas aeruginosa*, *Stenotrophomonas maltophilia* and *Mycobacteroides abscessus*. The observed overall selectivity of the hit compounds over the other homologues indicated structural differences between the proteins. We crystallized *M. smegmatis* and *P. aeruginosa* Arr enzymes, the former in complex with its most potent hit compound with an IC_50_ value of 1.3 µM. We observed structural differences in the NAD^+^ binding pockets of the two Arr homologues explaining the selectivity. Although the Arr inhibitors did not sensitize *M. smegmatis* to rifampicin in a growth inhibition assay, the structural information and the collection of inhibitors provide a foundation for rational modifications and further development of the compounds.

## Introduction

Rifampicin is a broad-spectrum antibiotic primarily used for the treatment of tuberculosis and other mycobacterial infections including leprosy and legionnaires’ disease (Hardie & Fenn, 2022). The bactericidal activity of rifampicin and other rifamycins derives from their binding to the beta subunit of bacterial RNA polymerase (RpoB) (**Figure 1A**), which blocks the elongation of the forming RNA transcript beyond 2-3 nucleotides.^1^ Although the predominant mechanism behind rifampicin resistance is point mutations in RpoB, for example in *Mycobacterium tuberculosis* most often on Ser531, His526 and Asp516,^2^ various bacterial species can also enzymatically inactivate the antibiotic, for instance, through ADP-ribosylation.^3^

**Figure 1.**
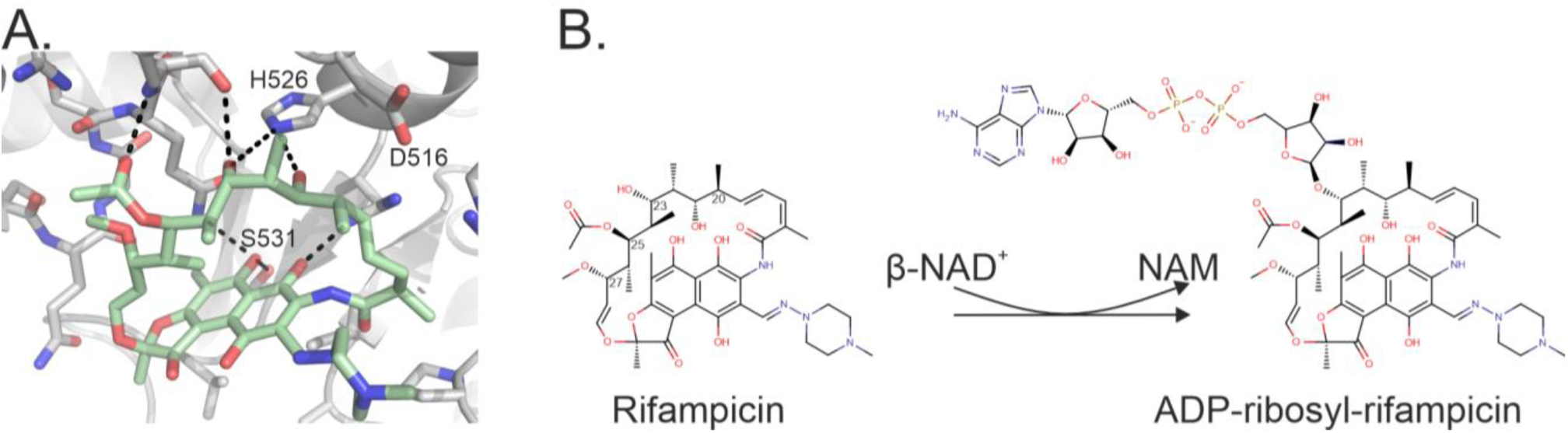
ADP-ribosylation of rifampicin inactivates its antibiotic effect. (A) Binding mode of rifampicin to the bacterial RNA polymerase of *Mycobacterium tuberculosis* (PDB id. 5UHB^7^). (B) ADP-ribosylation of rifampicin prevents its binding to the RNA polymerase as it occurs on the 23-hydroxyl group interacting with a key His526 in the binding site.

ADP-ribosylation has traditionally been recognized as a post-translational modification of proteins and ADP-ribosylation of nucleic acids has also gained attention recently.^4^ Currently, the only known small-molecule targets of ADP-ribosylation are rifamycin antibiotics. ADP-ribosylation of rifampicin was first identified in *Mycolicibacterium smegmatis,* whose Arr enzyme catalyzes the ADP-ribosylation of rifampicin and other rifamycin antibiotics.^5^ Arr transfers the ADP-ribose moiety from nicotinamide adenine dinucleotide (NAD^+^) to the 23-hydroxy group of rifampicin located in its ansa-bridge (**Figure 1B**). This leads to the inactivation of the antibiotic as it prevents its interaction with the beta subunit of RpoB.^6^

Although rifamycins are the only known targets of Arr-catalyzed ADP-ribosylation the protein has been linked to other catalytic and non-catalytic functions leading to a growth fitness advantage even in the absence of rifamycins.^8,9^ The expression of Arr is upregulated in response to various cellular stress stimuli and it has been associated with typical outcomes of the bacterial stringent response, including the accumulation of the guanosine tetraphosphate (ppGpp) alarmone, biofilm formation and decreased transcript levels of ribosomal proteins.^8^ Also, endogenous ADP-ribosylation of two unidentified proteins with molecular weights of 30 and 80 kDa have been detected in *M. smegmatis,* indicating that Arr likely has additional currently unknown cellular functions.^10^

The presence of Arr is not limited to *M. smegmatis.* It is widely spread among bacterial species being identified in 11 different phyla with high prevalence in Proteobacteria and Actinobacteria, especially in mycobacterial species.^11^ Arr has been found in both environmental and clinical isolates often associated with mobile genetic elements. One such example is Arr-2 originally identified from a clinical isolate of *P. aeruginosa* within a class 1 integron.^12^ Arr-2 has also been identified in several other pathogenic species, including *Klebsiella pneumoniae*^13^ and *Acinetobacter baumannii.*^14^ The presence of Arr in many pathogens combined with its apparent capacity for horizontal gene transfer and its role in antibiotic resistance indicate it could be a potential drug target.

Arr-mediated rifamycin resistance has previously been targeted with a natural rifamycin derivative kanglemycin A.^15^ Due to its bulky substituents in its ansa-bridge, specifically at positions C-20 and C-27, kanglemycin A cannot be modified by *M. smegmatis* Arr but retains its inhibitory effect on bacterial RNA polymerase. Additionally, several synthetic C-25 substituted rifamycins with improved activity against *M. smegmatis* and *M. abscessus* have been developed.^16–18^ A recent study showed that Arr-mediated rifamycin resistance could also be circumvented by directly inhibiting Arr enzymes.^19^ The authors discovered that AZ9482, an analogue of the PARP inhibitor Olaparib/Lynparza, inhibited *M. smegmatis* Arr with a 55 µM IC_50_ value in an HPLC-based activity assay. The compound seemingly sensitized *M. smegmatis* and *M. abscessus* to rifampicin, thereby validating the strategy to overcome Arr-mediated rifamycin resistance.

In this work, we developed a high-throughput assay to identify Arr inhibitors for multiple Arr homologues. The assay is homogeneous, cost-effective and the reagents are readily available. We applied the assay to screen an in-house library of human ADP-ribosyltransferase-targeted compounds and identified 15 compounds with IC_50_ values below 5 µM against four different Arr enzymes. The obtained co-crystal structure of MsArr in complex with the most potent hit compound confirmed its binding to the NAD^+^ binding site of the enzyme. The crystal structure of Arr-2, hereafter referred to as PaArr, revealed structural differences in the NAD^+^ binding pockets of the enzymes, which was also supported by the selectivity profile in the activity assays. The identified MsArr inhibitors did not sensitize *M. smegmatis* to rifampicin and we observed toxicity of the previously reported AZ9482 even in the absence of rifampicin. These results suggest that challenges affecting the activity of the compounds in cells, including impermeability of the mycobacterial cell wall and the efflux effect, must be addressed for MsArr to be a viable target. However, the assays developed in this work in combination with the obtained structural information will provide a foundation for further inhibitor development.

## Results and Discussion

### Assay development

To measure the activity of recombinantly produced *M. smegmatis* Arr we adapted a previously described activity assay.^20,21^ The assay is based on the quantification of NAD^+^ after the enzymatic reaction is stopped with the leftover NAD^+^ being chemically converted to a fluorophore. To optimize the protein activity and reproducibility of the signal on 384-well plates different buffer systems were tested (**Figure S1A-D**). Based on the optimization we selected 50 mM PIPES pH 7.5 containing 0.01% Triton X-100 as the buffer for all enzymatic assays. When testing the effect of different NAD^+^ and rifampicin concentrations, we observed that the maximal activity was reached when the rifampicin concentration was increased to twice the concentration of NAD^+^. On the other hand, excess of rifampicin inhibited the activity (**Figure 2A**). To achieve a robust signal during screening we decided to use the highest tested NAD^+^ concentration that did not lower its relative consumption and therefore selected 25 µM NAD^+^ (**Figure S1E**).

**Figure 2.**
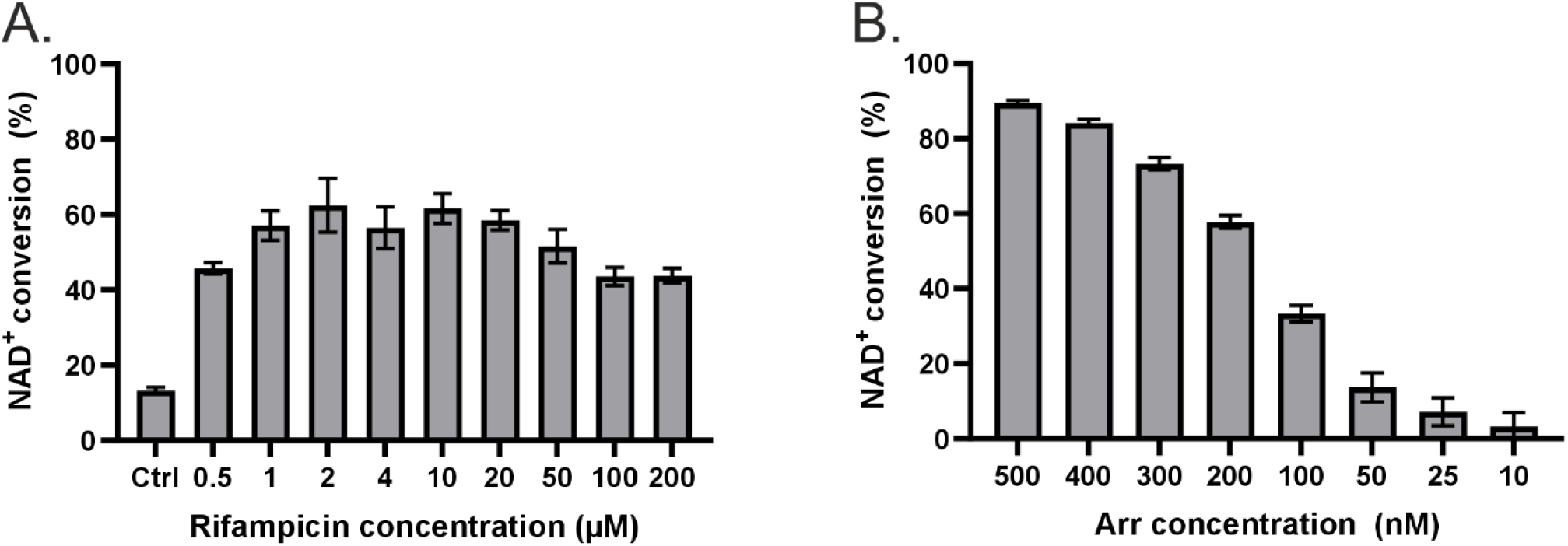
Activity tests with MsArr. (A) Effect of different rifampicin concentrations on the conversion of 1 µM NAD^+^ by 200 nM MsArr in 1 hour. (B) Testing the conversion of 1 µM NAD^+^ by different concentrations of MsArr in 1 hour.

Using the optimized conditions, we could reach 30–40% NAD^+^ consumption with 100 nM protein after 1 hour incubation, which is suitable for IC_50_ measurements (**Figure 2B**). For screening purposes, we aimed for 50-60% conversion to ensure robust assay performance, which was reached with 200-250 nM protein. If needed, the assay sensitivity could be improved by increasing the incubation time allowing lower protein concentration to be used as we have previously shown with human mono-ADP-ribosyltransferases.^22^ In a DMSO sensitivity test (**Figure S1F**) none of the tested concentrations up to 5% (v/v) DMSO affected the activity of MsArr and the highest concentration used in the assay was 1% (v/v). To assess the assay performance, we determined the signal-to-background ratio (S/B), signal-to-noise ratio, Z’-factor and coefficient of variation (CV) for minimal and maximum signals (**Table S1**). For this purpose, five plates were tested with a Z’-factor of 0.57 ± 0.03, indicating excellent assay quality.^23^

### Screening, hit validation and compound selectivity

We screened an in-house library of 538 human ADP-ribosyltransferase-targeted compounds against MsArr (**Figure 3A**). The library was screened in singlets at a concentration of 50 µM at which the reactions contained 0.5% (v/v) DMSO. We identified seven compounds with higher than 50% inhibition at this concentration. These hits were retested in dose-response experiments to exclude false positives and to obtain IC_50_ values (**Figure 3B**). The most potent ones, compound **1** and compound **2**, inhibited MsArr with IC_50_ values of 1.3 µM and 2.0 µM, respectively (**Figure 3B-C**).

**Figure 3.**
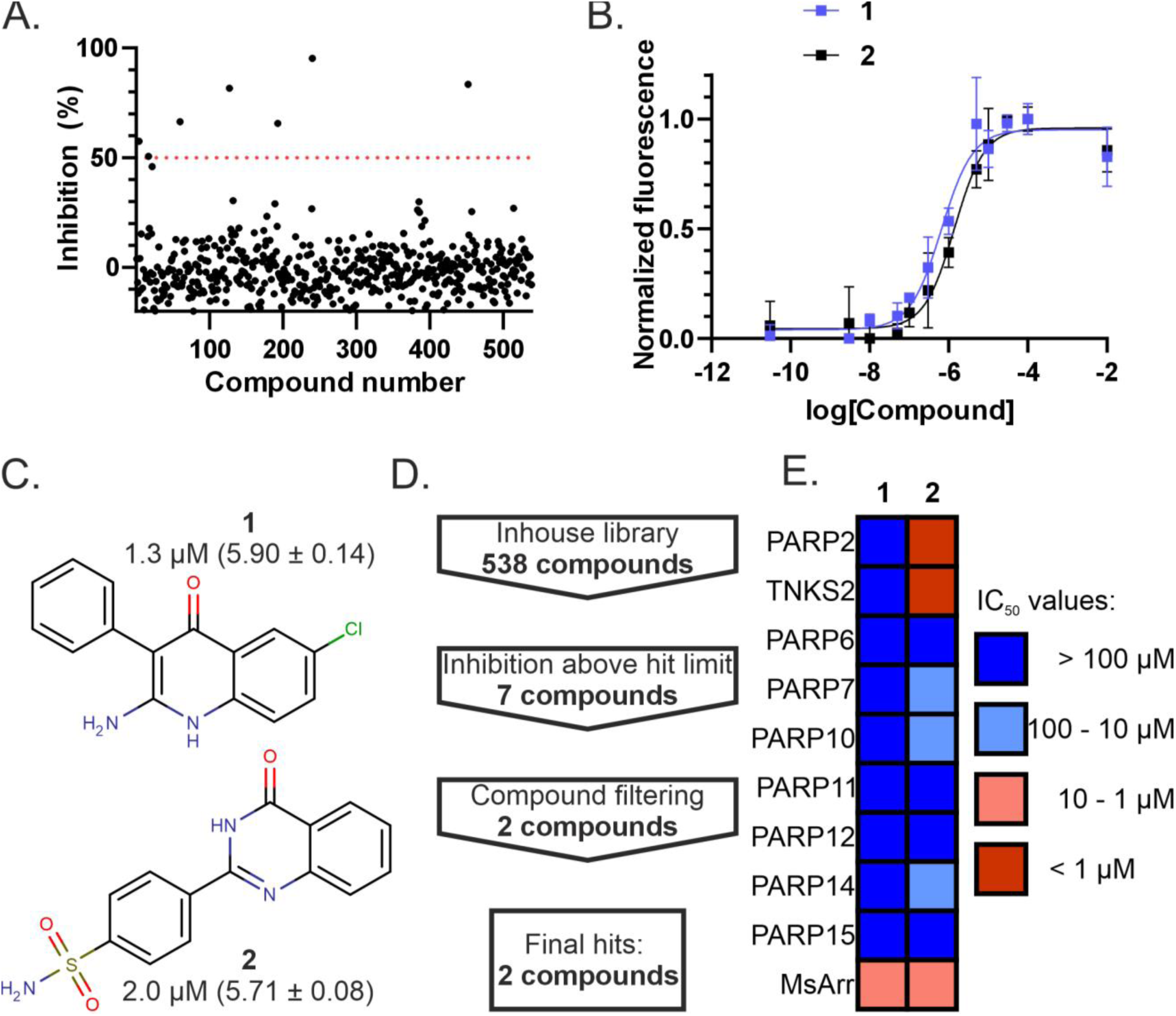
Screening of an in-house library against MsArr and evaluation of the hit compounds. (A) Results of the screening. (B) Examples of IC_50_ curves measured with the most potent compounds, **1** and **2**. (C) Structures and IC_50_ values (along with pIC_50_ and SEM, n=3) of **1** and **2**. (D) Workflow for identifying the hit compounds. (E) Heatmap of IC_50_ values of the compounds over a panel of human ADP-ribosyltransferases.

As the inhibition of Arr enzymes has obvious therapeutic potential we also wanted to check with a panel of enzymes to which extent the most potent compounds inhibit human ADP-ribosyltransferases. Compound **2** was previously reported to inhibit TNKS1 with an IC_50_ value of 56 nM^24^ and showed strong inhibition of TNKS2 and PARP2 as expected (**Figure 3E**), but did not inhibit the tested mono-ADP-ribosyltransferases. Compound **1**, which was originally synthesized to target human carbonic anhydrases^25^, did not inhibit any of the tested human ADP-ribosyltransferases (**Figure 3E**) making it a preferred compound for further development.

### Co-crystal structure of MsArr in complex with 1

We crystallized *M. smegmatis* Arr in complex with rifampicin and compound **1** (**Figure 4A**). The structure was solved with molecular replacement using a previously solved structure 2HW2 as a model and refined with data to 2.2 Å resolution resulting in R_work_ and R_free_ values of 0.19 and 0.25. Data collection and refinement statistics are shown in **Table S2**. The electron density for rifampicin is well-defined, covering the nicotinamide site as previously reported.^26^ Rifampicin contributes to the binding of **1** (**Figure 4B**) through a direct hydrogen bond between its 25-acetate group and the 2-amino group of the compound. Additionally, a jointly coordinated water molecule forms hydrogen bonds with both nitrogens of **1**, the side chains of Asp84 and Asn84 and the 23-hydroxy group of rifampicin.

**Figure 4.**
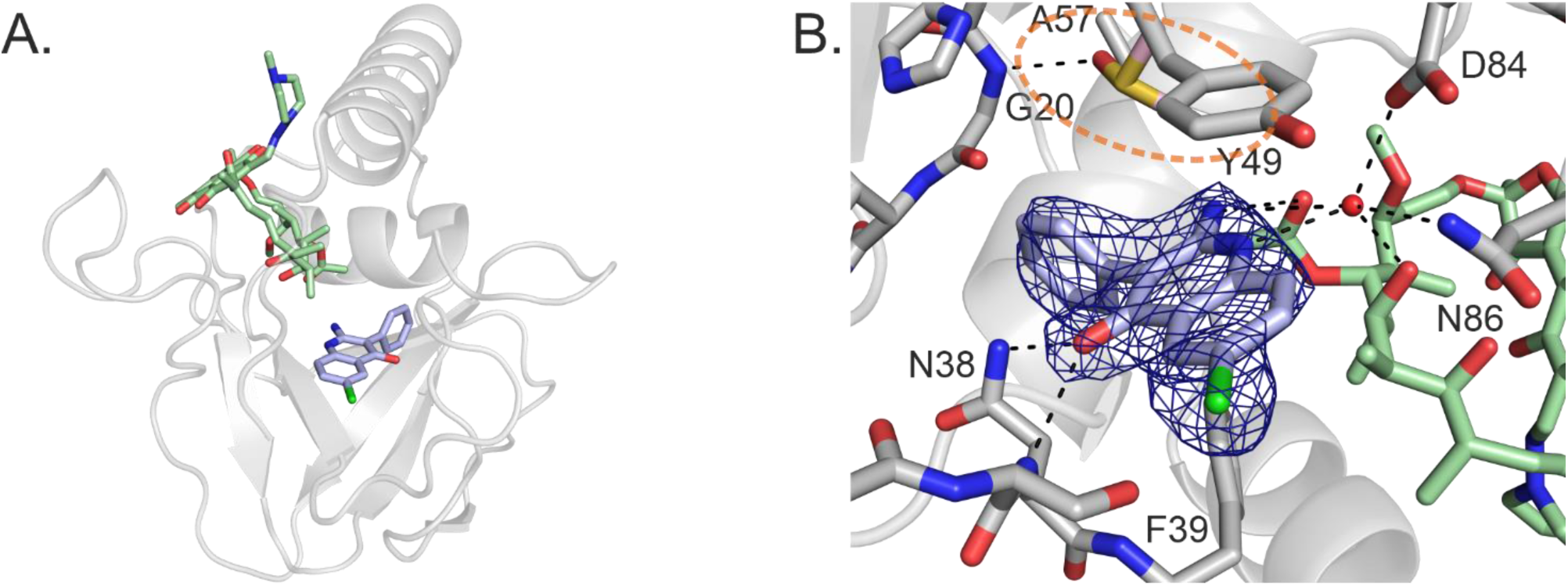
Co-crystal structure of MsArr (gray) with rifampicin (green) and **1** (blue). (A) Overall view of the MsArr structure with rifampicin and **1**. (B) A close-up view of the binding site of **1**. The nicotinamide binding site occupied by DMSO is indicated with an orange ellipse. The sigma A weighted 2Fo-Fc electron density map is presented in dark blue and contoured at 1 σ.

In contrast to most developed human ADP-ribosyltransferase inhibitors, **1** is not involved in hydrogen bonding with a serine side chain and glycine at the bottom of the nicotinamide pocket. In MsArr, the serine residue is replaced by Ala57 and, in our crystal structure, the backbone amide of Gly20 forms a hydrogen bond with DMSO instead (**Figure 4B**). The compound interacts with the MsArr loop Val32-His46 by hydrogen bonds between its carbonyl oxygen and the backbone amide and side chain of Asn38. Also, adjacent Phe39 is involved in T-shaped pi-stacking with the compound. Pi-stacking is also observed on the opposite side of the compound with Tyr49 (**Figure 4B**), which is also part of the catalytic H-Y-D motif and aligns with the tyrosine of the H-Y-E/I/L/Y catalytic motif of human ADP-ribosyltransferases. The loop Val32-His46 corresponds to the D-loop of human ADP-ribosyltransferases. However, its conformation differs from the typical position of the D-loops in crystal structures of human ADP-ribosyltranferases (**Figure S2**) offering an explanation for the observed selectivity over the tested human ADP-ribosyltranferases. The obtained co-crystal structure confirms the binding of the compound to the protein and allows rational design of improved analogues.

### *M. smegmatis* growth inhibition assay

After confirming that compound **1** binds to the protein and inhibits MsArr activity, we tested whether it could sensitize *M. smegmatis* to rifampicin. We measured the growth of the cells at three different rifampicin concentrations. Unfortunately, we did not observe any sensitization by **1** (**Figure 5A**). Although the quinazoline scaffold has been widely used in cell-based assays for tankyrase inhibitors, compound **2** did not show any signs of increased rifampicin susceptibility either.

**Figure 5.**
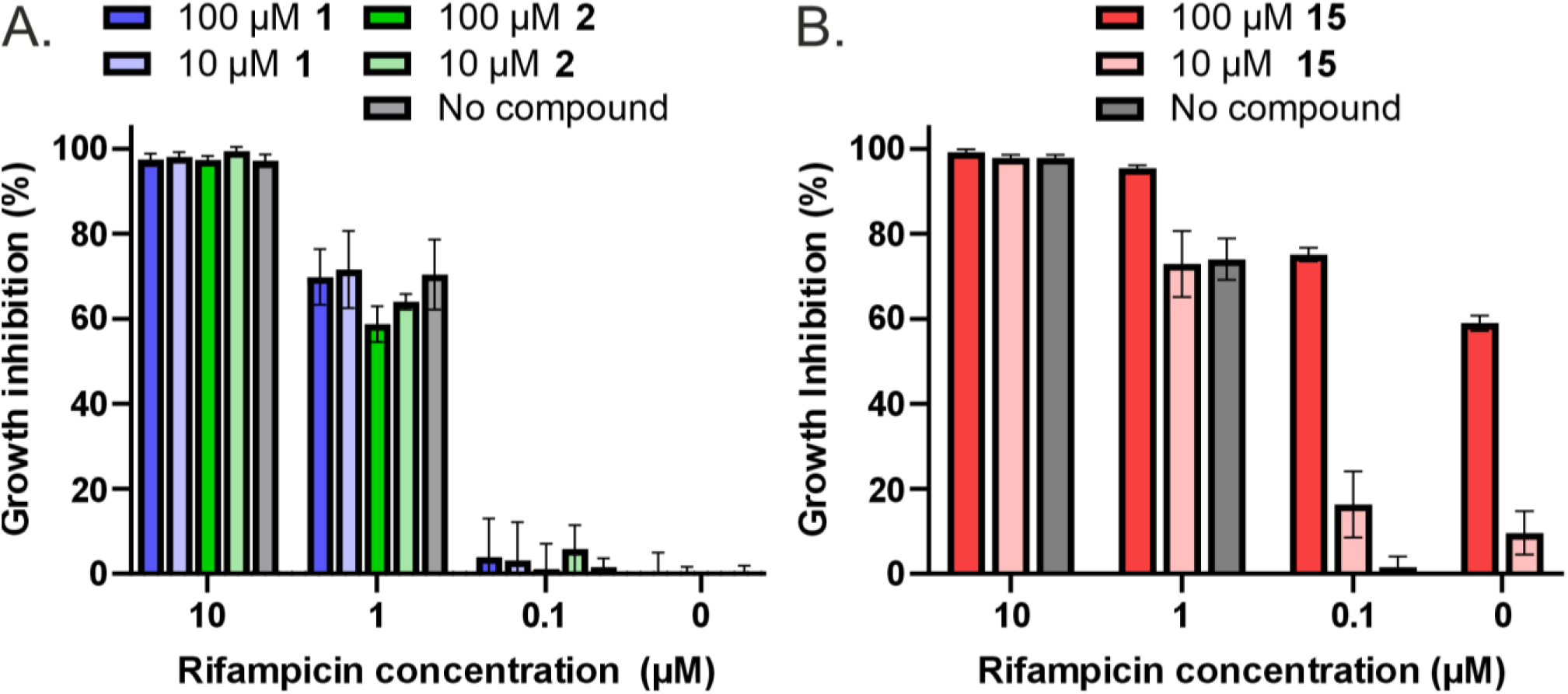
Proliferation of *M. smegmatis* in the presence of rifampicin and MsArr inhibitors. (A) Results with **1** and **2**. (B) Results with **15**. The reported values are mean inhibition percentages of three measurements.

Since our hit compounds failed to demonstrate activity in the cell assay, we wanted to confirm that, using our assay setup, we could reproduce previous results with **15**^19^, which was shown to inhibit MsArr and sensitize *M. smegmatis* and *M. abscessus* to rifampicin in a concentration-dependent manner. Using the developed NAD^+^ conversion assay, we measured an IC_50_ value of 35 µM, which is comparable to the lower potency hits, but 30 times less potent than compound **1**. In the cell assay, we were able to reproduce the results^19^ and observed a resembling inhibitory response (**Figure 5B**). However, it was evident that **15** inhibited cell growth even in the absence of rifampicin. Concentration-dependent growth inhibition reached 9.6% at 10 µM and was 59.0% at the highest tested concentration of 100 µM. Therefore, the observed growth inhibition in combination with rifampicin might not be related to the inhibition of Arr-catalyzed ADP-ribosylation of rifampicin.

The inability of the compounds to sensitize the cells to rifampicin might be explained by two primary factors. Mycobacteria are known for their lipid-rich cell wall, whose low permeability limits the uptake of antimicrobials.^27^ Additionally, several efflux pumps have been identified in *M. smegmatis*^28^ and their activity could prevent the compounds from reaching the target protein in cells. The combined effect of low permeability and active efflux could explain the lack of sensitization by the tested compounds, and they should be investigated in future studies.

### Identification of inhibitors of three additional Arr enzymes

We applied the developed activity assay to screen inhibitors for three additional Arr enzymes. The selected proteins were chosen based on their sequence divergence to understand how possible structural differences influence the inhibitor efficacy (**Figure S3**). PaArr, originally identified in *Pseudomonas aeruginosa* as Arr-2 in a class 1 integron,^12^ has later been identified in clinical isolates of several other rifampicin-resistant gram-negative pathogens.^13,14,29^ The second selected protein, SmArr, is specific to *Stenotrophomonas maltophilia,*^30,31^ another gram-negative pathogen where the gene was identified within a class 1 integron. MaArr, in turn, is a chromosomally encoded Arr enzyme from *Mycobacteroides abscessus*, previously described as Arr-Mab.^32^

We used the same conditions that were optimized earlier for MsArr and adjusted the protein concentrations to reach suitable conversion rates for screening and IC_50_ measurements with the additional proteins (**Figure S4A-C**). The final assay conditions for these proteins are listed in **Table S3**. The only exceptions were NAD^+^ and rifampicin concentrations which were lowered to 1 µM and 2 µM, as the relative NAD^+^ consumption of PaArr, SmArr and MaArr decreased at higher concentrations, likely due to NAD^+^ already being at saturated level (**Figure S4D**).

We screened the in-house library at a concentration of 25 µM. Even with the lower screening concentration the number of compounds exceeding the 50% inhibition limit was higher compared to MsArr for all three proteins. This was the case especially with PaArr, which was inhibited by 104 compounds. For SmArr and MaArr we observed 16 and 9 hits, respectively. The compounds interfering with the assay and the false positives were excluded again by IC_50_ measurements. The compounds with IC_50_ values below 5 µM are reported in **Figure 6**.

**Figure 6.**
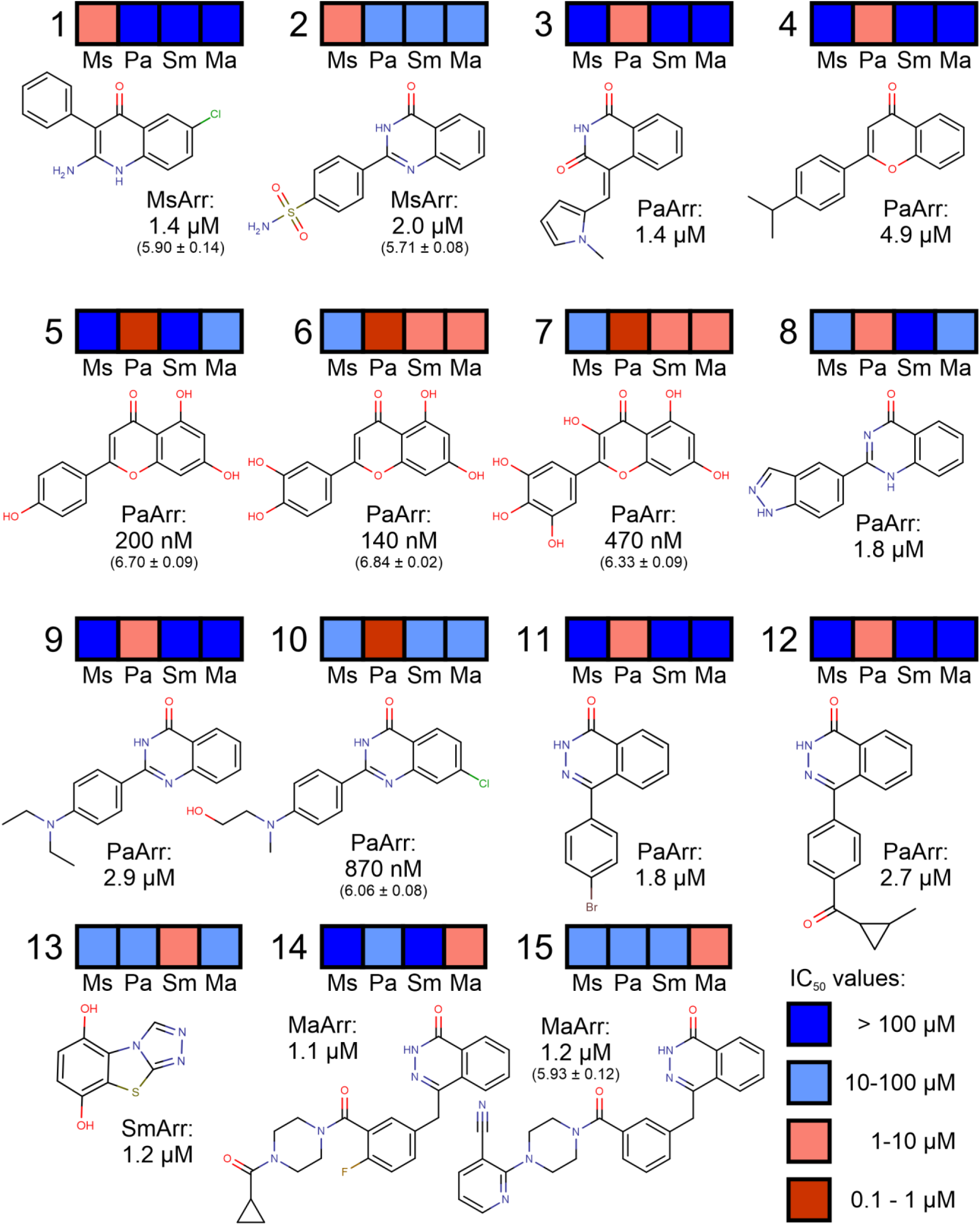
Final hit compounds against the four Arr enzymes and their selectivity profile. If the initial IC_50_ value was below 1 µM, the measurement was repeated three times and pIC_50_ and SEM values are reported.

We tested the inhibition of all four proteins with the 15 identified Arr inhibitors (**Figure 6**) and observed different inhibition profiles for the compounds (**Figure 6**). The MsArr inhibitor **1** was strongly selective over the other Arr enzymes showing less than 50% inhibition at 100 µM concentration against PaArr, SmArr and MaArr. In turn, with **2**, the IC_50_ values against the other enzymes fell between 10 and 100 µM. Out of the 12 compounds inhibiting PaArr, only compound **7** inhibited SmArr and MaArr with IC_50_ values below 10 µM. It was also the only compound inhibiting any other Arr enzyme with an IC_50_ value below the selected hit limit of 5 µM as it showed a 1.7 µM IC_50_ value against SmArr. The other SmArr inhibitor, **13**, was more selective as 50% inhibition was not reached at 10 µM compound concentration against the other enzymes. Interestingly, **15**, which we also tested in the *M. smegmatis* growth inhibition assays, was more potent as a MaArr inhibitor with an IC_50_ value of 1.2 µM showing 30-fold selectivity over MsArr. The diverse inhibition profiles of the compounds indicate structural differences among the Arr homologues.

While MsArr inhibitor **1** showed good selectivity over the tested human enzymes, the unselective compound **2** contains a known nicotinamide mimetic, quinazolinone, present in many PARP and tankyrase inhibitors. Known nicotinamide mimetic cores are also present in all PaArr, SmArr and MaArr hit compounds: isoquinolindione (**3**^33^, flavone (**4**-**7**^34,35^), quinazolinone (**8**-**10**^36^), triazolobenzothiazole (**13**^37^) or pthalazone (**11**-**12**, **14**-**15**).^38–40^. Many of these compounds inhibit human proteins at nanomolar concentrations, which should be considered if they are to be tested in infection models. Additionally, many human ARTDs are induced by infections, triggering cellular ADP-ribosylation, which should not be confused with the effects on rifampicin inactivation. The most suitable compound for such combination studies, based on our analysis, would be **1**, which lacks the nicotinamide mimicking moiety in contrast to the other hit compounds.

### Crystal structure of PaArr and comparison of Arr structures

Due to the lack of available structures and the observed selectivity, we attempted crystallization of PaArr. The protein crystallized as a monomer and the structure was solved at a resolution of 1.4 Å using a ColabFold^41^ prediction as a model for molecular replacement with R_work_ and R_free_ values of 0.14 and 0.18. Full data collection and refinement statistics are shown in **Table S2.** Although **10** was present in the crystallization conditions, we were unable to model it to the observed electron density in the NAD^+^ binding pocket, which was left unoccupied. The overall structure shows that the most prominent differences compared to the MsArr structure are seen in the C-terminal helix and the adjacent loop, which coordinates the binding of **1** in the case of MsArr (**Figure 7A**). In the crystal structure of PaArr, an additional rifampicin molecule stacks against these two areas. This causes the C-terminal half of the helix to be bent away from the additional rifampicin. The residues forming the loop surrounding the binding site of **1** in MsArr adopt a different conformation in PaArr, folding into strands β3 and β4, which are part of an additional β-sheet. This sheet is completed by a C-terminal strand, β8, originating from the bent helix. In contrast, the corresponding residues in MsArr occupy the region where the additional rifampicin binds in PaArr. The formed sheet acts as an extension to the strand β2 conserved in the ART fold and causes distinct differences in the shape of the NAD^+^ pocket compared to MsArr (**Figure 7B**). The additional strands protrude towards the nicotinamide site and overlap with the binding site of **1** explaining the observed lack of inhibition with PaArr. It should be noted that the observed additional rifampicin molecule is likely a crystallization artifact mediating crystal contacts and may not be biologically relevant. More importantly, its position and stacking against the areas surrounding the NAD^+^ binding pocket might affect the shape of the pocket. However, the observed selectivity profile where the identified MsArr inhibitors do not inhibit PaArr and vice versa supports the idea of substantial differences in the NAD^+^ binding pocket.

**Figure 7.**
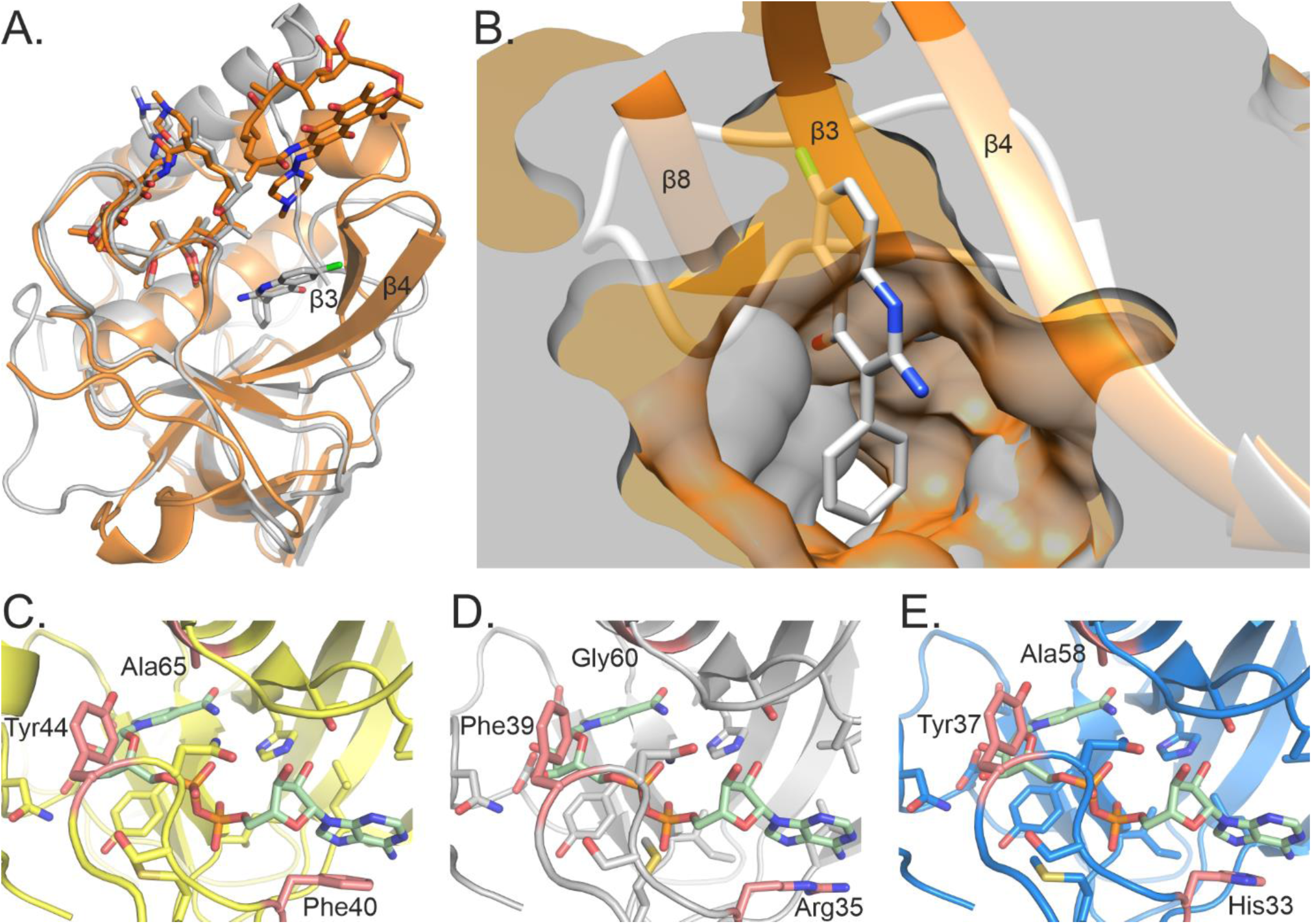
Superimposition of MsArr and PaArr structures and comparison of NAD^+^ binding pockets of different Arr enzymes. (A) Overall view of superimposed MsArr (gray) and PaArr (orange) including rifampicin molecules and **1**. (B) Superimposed protein surfaces highlighting compound **1** in the NAD^+^ pocket of MsArr. A loop surrounds the outer edge of the compound binding site in MsArr. In PaArr, the corresponding residues form additional β-strands that reshape the NAD^+^ binding pocket. The view is oriented from the rifampicin binding site towards the NAD^+^ pocket. (C) AlphaFold 3 prediction of *Stenotrophomonas maltophilia* Arr. (D) Our crystal structure of MsArr. (E) AlphaFold 3 prediction of *Mycobacteroides abscessus* Arr. NAD^+^ (green) is positioned in all three panels by deriving the coordinates from an AlphaFold 3 model of an MsArr-NAD^+^ complex. Residues surrounding the NAD^+^ pocket that differ between the three enzymes are highlighted in red.

The differently folded region not only binds NAD^+^ but is also involved in interactions with the acceptor molecules. As Arr has been reported to ADP-ribosylate protein substrates in addition to the evolutionary pressure to inactivate rifampicin, certain structural reorganizations of the region might have been necessary to efficiently target macromolecules and small molecules with different binding determinants. Previous structural studies have discovered only one conformation for the loop and therefore AlphaFold is guided towards predicting that one. However, our structure reveals that the loop can reorganize, which may have functional importance for binding different target molecules and partly explains the selectivity we observed in Arr inhibitors.

We compared the obtained MsArr crystal structure with AlphaFold 3 predictions (Abramson et al., 2024) of SmArr and MaArr to see if they could provide an explanation for the observed selectivity profile also in the case of these two proteins (**Figure 7C-E**). The overall folds of the AlphaFold predictions were identical to the MsArr crystal structure with minor differences in the orientation of the C-terminal helix of SmArr. Additionally, the residues surrounding the NAD^+^ binding site are conserved apart from three residues highlighted in **Figure 7C-E** that do not explain the selectivity if the hit compounds mimic the binding mode of nicotinamide. Therefore, there could be more drastic structural differences that AlphaFold cannot currently predict. This was already seen with PaArr, which was predicted to be folded identically to the other Arr enzymes including the loop that was folded into an additional β-sheet and reshaping the binding pocket.

In conclusion, we developed a high-throughput assay and utilized it to identify small-molecule inhibitors of multiple rifampicin resistance-conferring Arr enzymes. The results revealed structural differences in the NAD^+^ binding pockets of the homologues and demonstrate that issues related to the cellular efficacy of the inhibitors must be addressed if Arr enzymes are to be validated as viable drug targets. Nevertheless, Arr inhibition remains a promising strategy, and our work provides a framework for the rational design of more effective compounds to overcome rifamycin resistance of multiple pathogenic bacteria.

## Methods

### Cloning

MsArr V47I Y72F variant and the other Arr sequences were obtained as codon-optimized gBlock gene fragments (Integrated DNA Technologies) and cloned into pNIC-MBP vector^42^ by sequence and ligation-independent cloning (SLIC)^43^ in DH5α competent cells. Colonies were grown on LB agar containing 5% sucrose, utilizing the SacB-based negative selection marker.^44^ The constructs were subsequently verified by sequencing.

### Protein expression

The plasmids were transformed into *E. coli* BL21 (DE3) cells. Precultures were prepared in 5 ml of LB medium (Formedium) supplemented with 50 µg/ml kanamycin. After an 18-hour incubation at 37 °C, 500 ml of Terrific Broth (TB) autoinduction media including trace elements (Formedium), 8 g/l glycerol and 50 µg/ml kanamycin were inoculated with the preculture. The flasks were incubated at 37 °C until an OD_600_ of 0.6 was reached. The temperature was lowered to 18 °C and the incubation was continued overnight. Cells were harvested by centrifugation at 5020 g for 30 minutes at 4 °C. Finally, the cells were resuspended in lysis buffer (50 mM HEPES pH 7.5, 500 mM NaCl, 0.5 mM TCEP, 10 mM imidazole, 10% (v/v) glycerol) and frozen at −20 °C until purification.

### Protein purification

The cells resuspended in lysis buffer were sonicated in the presence of 0.05 mg/ml lysozyme, 0.02 mg/ml DNase I and 0.1 mM Pefabloc SC protease inhibitor (Roche) and the cell debris was removed by centrifugation at 18000 g for 40 minutes at 4°C. The supernatant was filtered through a syringe filter with a 0.45 µm pore size and then loaded onto a HiTrap Chelating HP IMAC column (Cytiva) pre-equilibrated in lysis buffer. The column was first washed with 5 column volumes of lysis buffer and then with 3-5 column volumes of wash buffer in which the imidazole concentration was increased to 25 mM. The protein was eluted with lysis buffer containing 350 mM imidazole. The MBP-tagged Arr proteins were passed over a 5 ml MBPTrap column (Cytiva) equilibrated with wash buffer and washed with 3 column volumes of the same buffer. The protein was eluted using the same buffer supplemented with 10 mM maltose. The His6-MBP-tag was cleaved overnight at 4°C by the addition of TEV protease (1:30 molar ratio) and removed by passing the sample over a HiTrap Chelating HP IMAC column equilibrated with wash buffer. The flowthrough containing MsArr was concentrated using a centrifugal filter with a 10 kDa MWCO (Amicon) and loaded onto a HiLoad 16/600 Superdex 75 pg (Cytiva) size-exclusion column equilibrated with 20 mM HEPES pH 7.5, 250 mM NaCl and 0.5 mM TCEP. Fractions containing the protein were combined, concentrated, aliquoted, flash-frozen in liquid nitrogen and stored at −70°C.

### NAD^+^ conversion Assay

Earlier described homogeneous activity assays for human PARP enzymes were adapted to measure the activity of MsArr.^20,21^ The assay is based on the quantification of leftover NAD^+^ after the enzymatic reaction is stopped and NAD^+^ is chemically converted to a flurophore. The assay was carried out on black round-bottom polypropylene 384-Shallow Well microplates (Thermo Fisher Scientific). Echo 650 acoustic liquid handler (Beckman Coulter) was used to dispense the tested compounds and DMSO to the assay plate while the other reagents were dispensed with Mantis liquid dispenser (Formulatrix). Final assay conditions are listed in **Table S3**. The reaction volume was 5 µl and DMSO was backfilled to a final concentration of 0.5% in screening and 1.0% in IC_50_ experiments. The assay plates were incubated for 1 hour at room temperature with shaking at 400 rpm on PST 100-HL plate incubator (Biosan).

The enzymatic reaction was stopped and the leftover NAD^+^ was converted to a fluorophore by the addition of 1 µl of 2 M KOH, 1 µl of 20 % acetophenone and 30 % glycerol in EtOH and 3 µl of formic acid, which was added 10 minutes after acetophenone. The fluorescent signal was measured with Infinite M1000 Pro plate reader (Tecan). The excitation and emission wavelengths were 372 nm and 444 nm, respectively.

### Protein crystallization and structure refinement

MsArr was co-crystallized with **1** and rifampicin using sitting-drop vapor diffusion method at room temperature. Protein and precipitant solutions were mixed in 1:1 ratio in 200 nl droplets using Mosquito crystallization robot (SPT Labtech). The droplets were monitored using with RI-54 imager (Formulatrix) through IceBear software.^45^ Protein solution contained 10 mg/ml MsArr (630 µM), 1 mM **1** and 1 mM rifampicin. The protein aliquot was diluted at a 1:1 ratio in water when preparing the solution. The precipitant solution contained 50 mM Tris pH 8.2, 0.2 M MgCl_2_, 10% (w/v) PEG 8000, 20% (v/v) 2-methyl-2,4-pentanediol (MPD). The crystals were cryoprotected with the precipitant solution which was mixed with glycerol in a MPD in a 7:3 ratio.

PaArr was co-crystallized with **10** in a similar way as MsArr using protein-precipitant ratio of 1:2 in a 225 nl droplet. The protein solution contained 5 mg/ml PaArr (293 µM), 1 mM **10** and 1 mM rifampicin. The protein aliquot was diluted at a 1:3.16 ratio in water. The precipitant solution contained 100 mM MOPS pH 7.5, 20% ethanol, and 2.7 M (NH_4_)_2_ SO_4_. The crystals were cryoprotected with the precipitant solution which was mixed with glycerol in a 4:1 ratio.

Collected datasets were processed with XDS.^46^ Molecular replacement was done using MOLREP^47^ in CCP4i2 using PDB id 2HW2^26^ as a search model for MsArr and ColabFold^41^ prediction for PaArr. The model was built in Coot^48^ and refinement was done using Refmac5.^49^ Structure images were created using PyMol (The PyMOL Molecular Graphics System, Version 2.5.0, Schrödinger, LLC) and Chimera.^50^ Structural predictions of SmArr and MaArr were generated with AlphaFold.^51^

### *M. Smegmatis* growth inhibition assay

*M. smegmatis* MC^2^ 155 cultures were grown at 37 degrees in Middlebrook 7H9 broth (Sigma) containing 10 % ADS, 0.05 % Tween-80 and 0.2 % glycerol. The cultures were diluted from mid-logarithmic phase to OD_600_ value of 0.01 twice before the MIC_50_ measurements were started. The MIC_50_ measurements were conducted in 50 ml FastGene centrifuge tubes (Nippon Genetics) using a culture volume of 2 ml. DMSO concentration in each sample was backfilled to 1.1% (v/v). The cells were grown for 24 hours and their OD_600_ values were measured. The absorbances were measured with Ultrospec 10 cell density meter (Biochrom).

## Author contributions

Conceptualization: STS

Data curation: JA, LL

Formal analysis: JA, SA, STS, LL

Funding acquisition: LL

Methodology: JA, SA, STS

Investigation: JA, SA, STS

Project administration: LL

Resources: LL

Visualization: JA

Validation: JA, LL

Supervision: LL

Writing—original draft: JA

Writing—review & editing: JA, SA, STS, LL

The authors have no competing interests.

## Data availability

Atomic coordinates and structure factors will be available at the Protein Data Bank with identifiers 9IAF and 9IBA. Raw diffraction data will be available at fairdata.fi (https://doi.org/10.23729/a14040e9-f046-42c5-b7ab-7d9c53dc0bae).

## Acknowledgements

The use of the facilities and expertise of the Biocenter Oulu Structural Biology Core Facility, a member of Biocenter Finland, Instruct-ERIC Centre Finland and FINStruct, are gratefully acknowledged. We thank the local contacts for their assistance in using beamlines ID30A-1 at ESRF and I04 at Diamond Light Source. We thank Johan Pääkkönen for the help with crystallography. Rajaram Venkatesan, Nora Tir-Hynönen and Mikko J. Hynönen are thanked for the support in using the Biosafety level 2 laboratory. We thank Gianfranco Balboni for providing compound **1**. We acknowledge all our collaborators who have contributed to the in-house targeted inhibitor library over the years.

