## Supplementary information for "Discovery of inhibitors for bacterial Arr enzymes ADP-ribosylating and inactivating rifamycin antibiotics"

**CONTENT**

**Figure S1.** Optimization of the MsArr assay conditions.

**Table S1.** Assay performance.

**Table S2.** Data collection and refinement statistics.

**Figure S2.** Structural alignment of MsArr with human ADP-ribosyltransferases.

**Figure S3.** Sequence comparison of the different Arr homologs.

**Figure S4.** Assay development with PaArr, SmArr and MaArr.

**Table S3.** Assay conditions for the four Arr enzymes

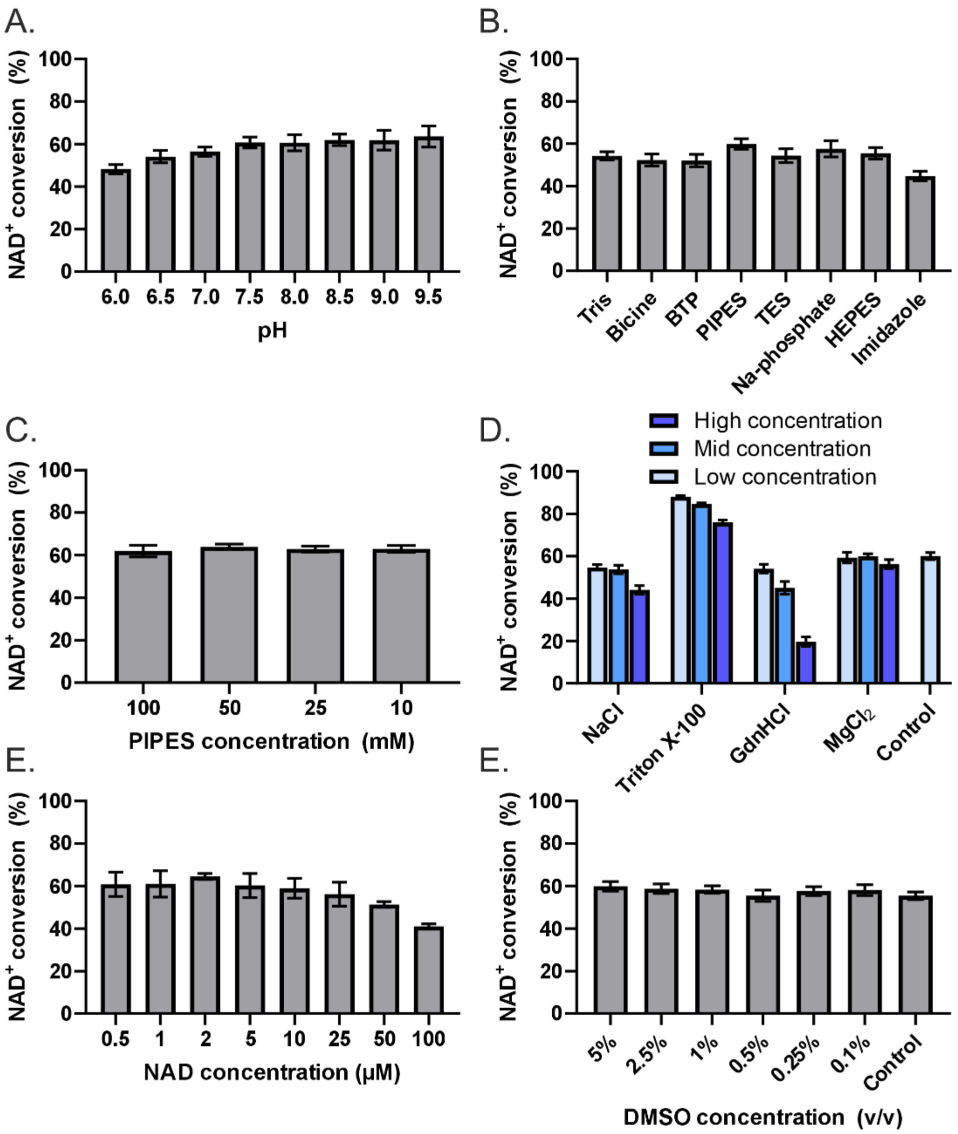


**Figure S1.** Optimization of the MsArr assay conditions. (A) Buffer pH optimization in 50 mM Bis-TRIS-propane. (B) Buffering reagent optimization in 50 mM buffer at pH 7.5. (C) Optimization of PIPES concentration at pH 7.5. (D) Effect of selected buffer components in 50 mM PIPES at pH 7.5. Used concentrations for the components were: 500 mM, 150 mM and 50 mM for NaCl, 0.25%, 0.05% and 0.01% (v/v) for Triton X-100, 500 mM, 150 mM and 50 mM, 10 mM for GdnHCl and 10 mM, 2.5 mM and 0.1 mM for $MgCl_2$ E) Testing different NAD and rifampicin concentrations using 150 nM protein and constant rifampicin/$NAD^+$ ratio of 2. (F) DMSO sensitivity test using 150 nM protein. Incubation time is 1 hour in all experiments. $NAD^+$ and protein concentrations are 1 μM and 500 nM, respectively, except in (E) where $NAD^+$ concentration was varied. Data shown as mean ± SD of 4 replicates.

**Table S1.** Assay performance.

| | **MsArr** | **PaArr** | **SmArr** | **MaArr** |
|---|---|---|---|---|
| S/B | 2.7 ± 0.1 | 2.3 ± 0.1 | 3.7 ± 0.5 | 2.7 ± 0.1 |
| S/N | 9.9 ± 0.8 | 11 ± 0.8 | 9.9 ± 2.2 | 21 ± 2.1 |
| *Z′* | 0.57 ± 0.03 | 0.57 ± 0.08 | 0.60 ± 0.09 | 0.80 ± 0.02 |
| Plate-to-plate, CV (%) | 7.5 | 15 | 19 | 3.9 |
| Day-to-day, CV (%) | 8.0 | 15 | 15 | 3.2 |

**Table S2.** Data collection and refinement statistics.

| | MsArr (PDB id. 9IAF) | PaArr (PDB id. 9IBA) |
|---|---|---|
| **Data collection** | | |
| Beamline | I04 | ID30A-1 |
| Wavelength (Å) | 0.953745 | 0.96546 |
| Space group | C2 | $P2_12_12_1$ |
| Cell dimensions | | |
| *a*, *b*, *c* (Å) | 56.2, 60.8, 45.6 | 41.3, 43.0, 85.5 |
| α, β, γ (°) | 90.0. 93.5, 90.0 | 90, 90, 90 |
| Resolution (Å) | 50 - 2.20 | 50 – 1.40 |
| Outer shell (Å) | 2.20 - 2.26 | 1.40 - 1.44 |
| No. unique reflections | 7819 (569) | 30653 (2251) |
| $R_{merge}$ | 0.120 (1.030) | 0.066 (0.802) |
| Mean $I/\sigma I$ | 12.53 (1.87) | 12.41 (2.12) |
| CC ½ (%) | 99.9 (79.7) | 99.8 (78.4) |
| Completeness (%) | 99.4 (96.4) | 99.6 (99.8) |
| Redundancy | 11.0 | 5.03 (5.12) |
| **Refinement** | | |
| $R_{work}$ / $R_{free}$ | 0.187 / 0.250 | 0.143 / 0.185 |
| No. atoms | | |
| Protein | 1099 | 1168 |
| Ligand/ion | 82 | 118 |
| Water | 28 | 115 |
| *B*-factors | | |
| Protein | 48.73 | 22.96 |
| Ligand/ion | 54.51 | 21.92 |
| Water | 43.26 | 31.12 |
| R.m.s. deviations | | |
| Bond lengths (Å) | 0.0124 | 0.0116 |
| Bond angles (°) | 1.865 | 1.662 |
| Ramachandran plot (%) | | |
| Favored | 133 (96.4%) | 141 (98.6%) |
| Allowed | 5 (3.6%) | 1 (0.7%) |
| Outliers | 0 | 1 (0.7%) |

Values in parentheses are for the highest-resolution shell.

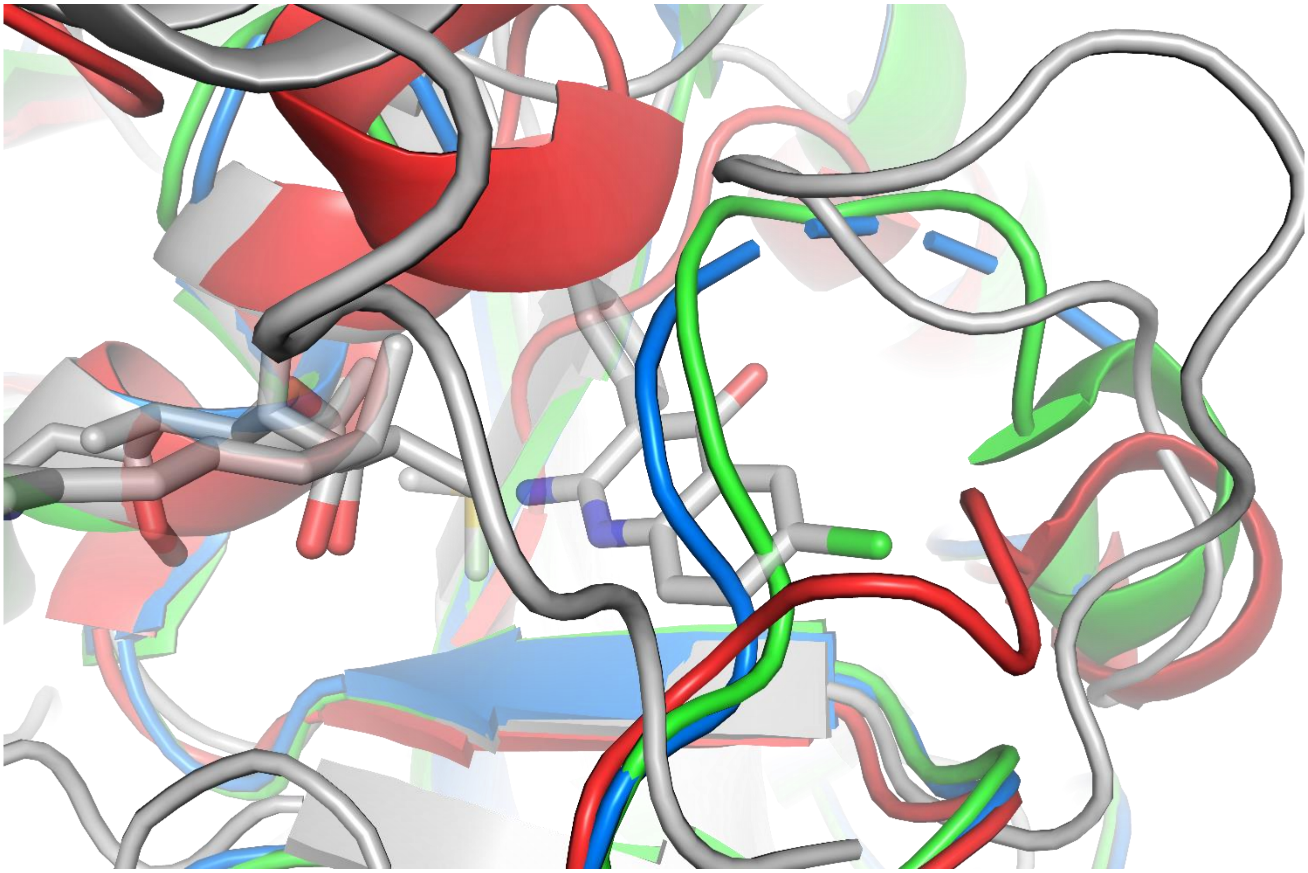

**Figure S2.** Structural alignment of MsArr Val32-His46 loop with D-loops of selected human ADP-ribosyltransferases. PARP10 is shown in green (PDB id. 6FXI), PARP15 is shown in blue (PDB id. 7Z2Q[1]) and TNKS2 is shown in red (PDB id. 7O6X[2]).

A.

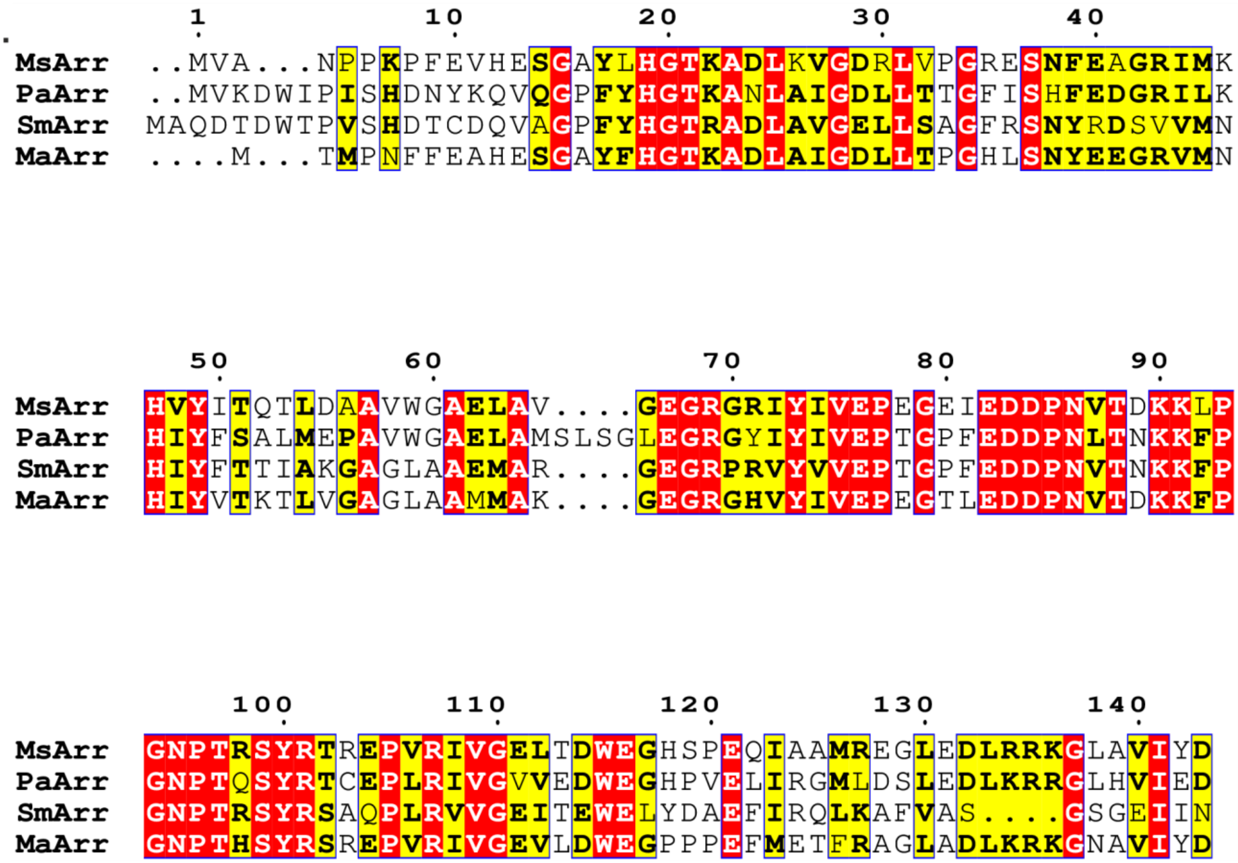


B.

| | MsArr | PaArr | SmArr | MaArr |
|---|---|---|---|---|
| MsArr | 100 | 57.34 | 43.88 | 65.96 |
| PaArr | 57.34 | 100 | 52.11 | 52.48 |
| SmArr | 43.88 | 52.11 | 100 | 52.55 |
| MaArr | 65.96 | 52.48 | 52.55 | 100 |

**Figure S3.** Sequence comparison of the different Arr homologs. (A) Multiple sequence aligment of the Arr homologs made with Clustal Omega[3] and presented with Espript 3.0.[4] (B) Sequence identity matrix of the Arr homologs (%).

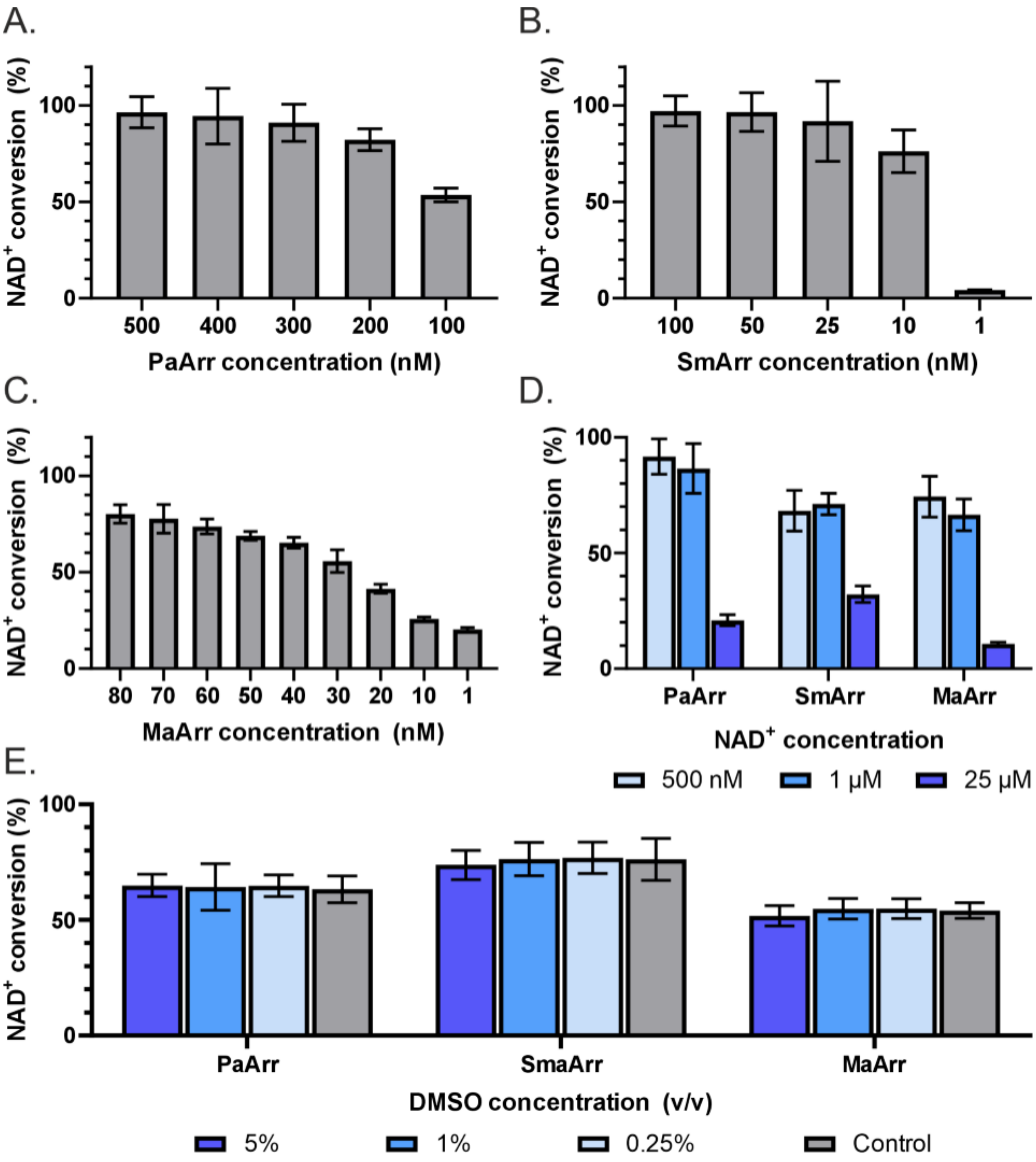


**Figure S4.** Assay development with PaArr, SmArr and MaArr. (A-C) Testing different protein concentrations. (D) Testing different $NAD^+$ concentrations in final assay conditions with constant Rifampicin/$NAD^+$ ratios of 2. (E) DMSO sensitivity test. Data shown as mean ± SD of 4 replicates.

**Table S3.** Final assay conditions for the four Arr enzymes

| | MsArr | PaArr | SmArr | MaArr |
|---|---|---|---|---|
| Buffer | 50 mM PIPES pH 7.5 0.01 % Triton X-100 | 50 mM PIPES pH 7.5, 0.01 % Triton X-100 | 50 mM PIPES pH 7.5, 0.01 % Triton X-100 | 50 mM PIPES pH 7.5, 0.01 % Triton X-100 |
| $NAD^+$ concentration | 1 µM ($IC_{50}$ assay) 25 µM (screening) | 1 µM | 1 µM | 1 µM |
| Rifampicin concentration | 2 µM ($IC_{50}$ assay) 50 µM (screening) | 2 µM | 2 µM | 2 µM |
| Protein concentration | 100 nM ($IC_{50}$ assay) 200 nM (screening) | 90 nM ($IC_{50}$ assay) 110 nM (screening) | 5.5 nM $IC_{50}$ assay) 6.5 nM (screening) | 20 nM ($IC_{50}$ assay) 30 nM (screening) |
| Incubation time | 1 hour | 1 hour | 1 hour | 1 hour |

(2) Leenders, R. G. G.; Brinch, S. A.; Sowa, S. T.; Amundsen-Isaksen, E.; Galera-Prat, A.; Murthy, S.; Aertssen, S.; Smits, J. N.; Nieczypor, P.; Damen, E.; Wegert, A.; Nazaré, M.; Lehtiö, L.; Waaler, J.; Krauss, S. Development of a 1,2,4-Triazole-Based Lead Tankyrase Inhibitor: Part

II. *J Med Chem* **2021**, *64* (24), 17936–17949. https://doi.org/10.1021/acs.jmedchem.1c01264.
(3) Madeira, F.; Madhusoodanan, N.; Lee, J.; Eusebi, A.; Niewielska, A.; Tivey, A. R. N.; Lopez, R.; Butcher, S. The EMBL-EBI Job Dispatcher Sequence Analysis Tools Framework in 2024. *Nucleic Acids Res* **2024**, *52* (W1), W521–W525. https://doi.org/10.1093/nar/gkae241.
(4) Robert, X.; Gouet, P. Deciphering Key Features in Protein Structures with the New ENDscript Server. *Nucleic Acids Res* **2014**, *42* (Web Server issue), W320-324. https://doi.org/10.1093/nar/gku316.
